# Origin and evolution of the cannabinoid oxidocyclase gene family

**DOI:** 10.1101/2020.12.18.423406

**Authors:** Robin van Velzen, M. Eric Schranz

**Author notes:** Author for Correspondence: Robin van Velzen, Biosystematics Group, Wageningen University, Wageningen, The Netherlands.

## Abstract

*Cannabis* is an ancient crop representing a rapidly increasing legal market, especially for medicinal purposes. Medicinal and psychoactive effects of *Cannabis* rely on specific terpenophenolic ligands named cannabinoids. Recent whole-genome sequencing efforts have uncovered variation in multiple genes encoding the final steps in cannabinoid biosynthesis. However, the origin, evolution, and phylogenetic relationships of these cannabinoid oxidocyclase genes remain unclear. To elucidate these aspects we performed comparative genomic analyses of *Cannabis*, related genera within the Cannabaceae family, and selected outgroup species. Results show that cannabinoid oxidocyclase genes originated in the *Cannabis* lineage from within a larger gene expansion in the Cannabaceae family. Localization and divergence of oxidocyclase genes in the *Cannabis* genome revealed two main syntenic blocks, each comprising tandemly repeated cannabinoid oxidocyclase genes. By comparing these blocks with those in genomes from closely related species we propose an evolutionary model for the origin, neofunctionalization, duplication, and diversification of cannabinoid oxidocycloase genes. Based on phylogenetic meta-analyses, we propose a comprehensive classification of three main clades and seven subclades that is intended to aid unequivocal referencing and identification of cannabinoid oxidocyclase genes. Our data suggest that cannabinoid oxidocyclase gene copy number variation may have less functional relevance than previously thought. Instead, we propose that cannabinoid phenotype is primarily determined by presence/absence of single-copy genes. Increased sampling across *Cannabis’* native geographic range is likely to uncover additional cannabinoid oxidocyclase gene sequence variation.

**Significance statement:** *Cannabis* genome sequencing efforts have revealed extensive cannabinoid oxidocyclase gene variation. However, phylogenetic relationships and evolution of these genes remains unclear. Our meta analysis of currently available data reveals that these genes comprise three main clades and seven subclades that originated through *Cannabis*-specific gene duplication and divergence. Our new conceptual and evolutionary framework serves as a reference for future description and functional analyses of cannabinoid oxidocyclases.

## Introduction

The plant *Cannabis sativa* L. (henceforth *Cannabis*) is an ancient yet controversial crop. *Cannabis* cultivars are commonly divided into “Fiber-type” (or hemp) cultivars that are used for the production of fibre and/or seed oil and “drug-type” (or marijuana) cultivars that are used for recreational, ritual, or medicinal purposes (Small and Cronquist 1976; McPartland and Guy 2017). Widespread recreational (ab)use led to its classification as a most highly restricted “schedule IV” class controlled substance (United Nations General Assembly 1975). However, recent scientific investigations have confirmed its medicinal benefits for various patients and ailments (Abrams 2018). Accordingly, the United Nations Commission on Narcotic Drugs recently voted to reclassify *Cannabis* as the less-stringent “schedule I” class and many countries are now legalizing some of its uses (Anon 2020; Hammond et al. 2020).

Consequently, *Cannabis* currently represents a rapidly emerging legal industry with an estimated multi-billion global market, primarily for medicinal purposes. However, fundamental aspects about the molecular evolution of *Cannabis* remain unknown (Hurgobin et al. 2020; Kovalchuk et al. 2020). In this paper, we aim to elucidate the origin and evolution of a unique class of biosynthetic genes found in the *Cannabis* genome.

Many of the medicinal properties of *Cannabis* are due to its production of cannabinoids; a unique class of psychoactive terpenophenolic ligands (Gaoni and Mechoulam 1964; Mechoulam 2005). Cannabinoids interfere with the human central nervous system through the endocannabinoid receptors which exist in the mammalian brain and immune cells (Pertwee 2008; Mechoulam and Parker 2013). The two most abundant and well-known cannabinoids are Δ^9^-tetrahydrocannabinol (THC) and cannabidiol (CBD), but more than 120 others have been identified in *Cannabis* (ElSohly et al. 2017). Some other plant genera such as *Rhododendron* and *Radula* have also been found to make cannabinoids (Iijima et al. 2017; Gülck and Møller 2020). THC is responsible for the psychoactive effect of *Cannabis* through its partial agonist activity at endocannabinoid receptors (Gaoni and Mechoulam 1964; Mechoulam and Parker 2013). This effect is the reason for the large-scale use of *Cannabis* as an intoxicant. But accumulating evidence from clinical trials indicate that moderate doses of THC can be used medicinally to e.g. reduce nausea and vomiting, pain, and improvement of sleep and appetite (van de Donk et al. 2019; Grimison et al. 2020; Suraev et al. 2020). CBD has weak affinity for endocannabinoid receptors and is not psychoactive (Pertwee 2005). It has been found to modulate the effects of THC and endocannabinoids and may be effective for symptomatic treatment of anxiety and psychosis and in treating some childhood epilepsy syndromes (Gofshteyn et al. 2017; Bhattacharyya et al. 2018; Skelley et al. 2020).

Fiber-type cultivars typically have low content (<0.4%) of THC and intermediate content (2-4%) of CBD while drug-type cultivars typically have high content (14-40%) of THC and low content (<1%) of CBD (Small and Cronquist 1976). However, cultivars exist with alternative chemical profiles such as drug-type cultivars with high levels of CBD and other classifications based on chemotype have been proposed (Hazekamp et al. 2016; Wenger et al. 2020). Current breeding programmes aim to develop new cultivars with specific cannabinoid profiles tuned for specific medicinal effects or consumer preferences. Some *Cannabis* plants synthesize cannabichromene (CBC) or cannabigerol (CBG) (Morimoto et al. 1997; Fournier et al. 2004). These lesser-known cannabinoids may have anti-inflammatory effects but evidence is relatively scarce (Brierley et al. 2019; Udoh et al. 2019). It is important to note that cannabinoids are synthesised and stored in the plant as acids that are not medicinally active. Only from exposure to light during storage or heat during processing for consumption (e.g smoking or baking) these acids are non-enzymatically decarboxylated to their neutral forms that have psychoactive and/or medicinal properties.

### The cannabinoid biosynthetic pathway

Within the *Cannabis* plant, cannabinoids are synthesized in multicellular epidermal glands (glandular trichomes) that are most abundant on the bracts of female inflorescences. Glandular trichomes are composed of a stalk and a gland, the latter comprising a set of secretory cells and a single large apoplastic storage cavity. The cannabinoid biosynthetic pathway has been largely elucidated, and for many steps in the pathway the corresponding enzymes have been isolated and characterized (Figure 1). In brief, cannabinoid biosynthesis relies on two precursors from two distinct metabolic pathways: olivetolic acid from the polyketide pathway and geranyl-geranyl pyrophosphate (GPP) from the methylerythritol phosphate (MEP) pathway. Biosynthesis of olivetolic acid occurs in the cytosol and starts with activation of hexanoic acid by acyl activating enzyme 1 into hexanoyl-CoA (Stout et al. 2012). Next, aldol condensation of hexanoyl-CoA with three molecules of malonyl-CoA leads to olivetolic acid. This reaction requires both the polyketide synthase named olivetol synthase (OLS) and the DABB-type polyketide cyclase named olivetolic acid cyclase (OAC) (Raharjo et al. 2004; Taura et al. 2009; Gagne et al. 2012). Olivetolic acid is then transported via an unknown mechanism to the plastid where it converted into cannabigerolic acid (CBGA) by c-terminal prenylated with GPP by CBGAS - a transmembrane aromatic prenyltransferase with a plastid localization signal (Fellermeier and Zenk 1998; Luo et al. 2019). CBGA is then secreted into the extracellular storage cavity via an unknown mechanism and further processed by secreted cannabinoid oxidocyclases that perform different types of oxidative cyclizations of its linear prenyl moiety into derived cannabinoid acids such as tetrahydrocannabinolic acid (THCA), cannabidiolic acid (CBDA), and cannabichromenic acid (CBCA) (Taura et al. 1995; Taura et al. 1996; Morimoto et al. 1998; Sirikantaramas et al. 2005; Rodziewicz et al. 2019). *Cannabis* plants accumulating CBGA are assumed to have nonfunctional cannabinoid oxidocyclases (De Meijer and Hammond 2005; Onofri et al. 2015).

**Fig 1.**
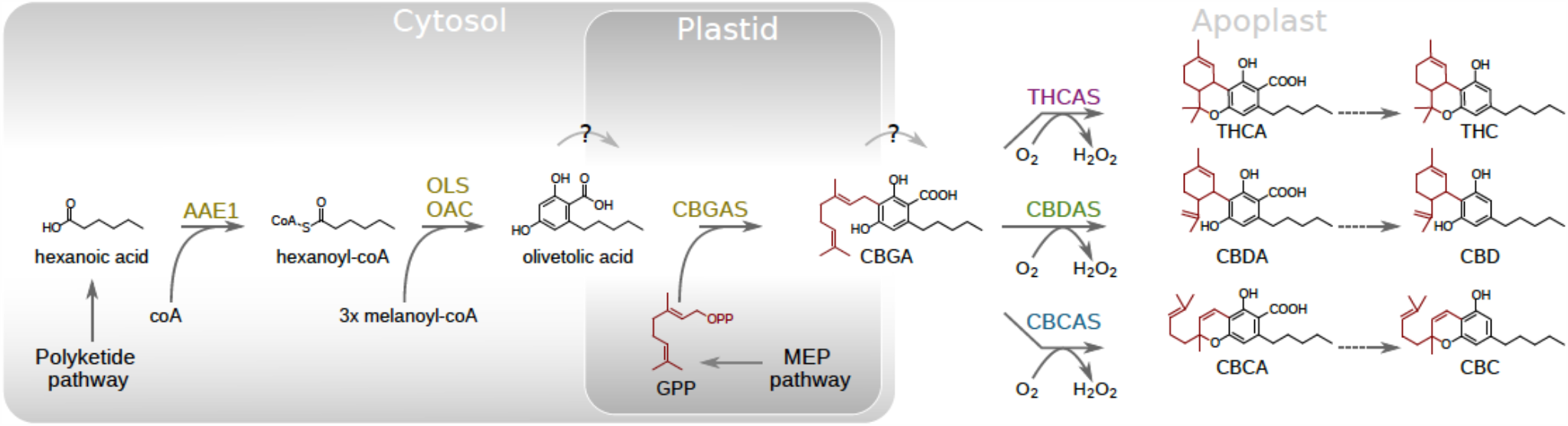
Cannabinoid biosynthesis pathway. Dotted arrows indicate non-enzymatic decarboxylations; solid arrows indicate enzymatic reactions; Enzyme names are shown in blue, while resulting compounds are shown in black. Compound (sub)structures depicted in red signify those that represent a single unit of GPP. Abbreviations: AAE1, acyl-activating enzyme 1; CBC, cannabichromene; CBCA(S),cannabichromenic acid (synthase); CBD, cannabidiol; CBDA(S), cannabidiolic acid (synthase); CBGA(S),cannabigerolic acid (synthase); GPP(S), geranyl-pyrophosphate (synthase); MEP, methylerythritol phosphate; OAC, olivetolic acid cyclase; OLS, olivetol synthase; THC, tetrahydrocannabinol; THCA(S), tetrahydrocannabinolic acid (synthase).

The three currently known cannabinoid oxidocyclase enzymes THCA synthase (THCAS), CBDA synthase (CBDAS), and CBC synthase (CBCAS) are highly similar in their biochemical properties and sequence characteristics (Taura, Sirikantaramas, Shoyama, Shoyama, et al. 2007; Laverty et al. 2019). The masses of the mature proteins are estimated to be 76 kDa for THCAS, 74 kDa for CBDAS, and 59 kDa for CBCAS. In terms of catalytic kinetics, THCAS and CBDAS have highly similar affinities to CBGAS as a substrate (K_m_ = 137 and 134 μM, respectively) and turnover number (K_cat_ = 0.20 and 0.19 s^-1^, respectively) (Taura et al. 1995; Taura et al. 1996). In comparison, CBCAS has a higher substrate affinity (K_m_ = 23 μM) and lower catalytic rate constant (k_cat_ = 0.04 s^-1^) (Morimoto et al. 1998; Laverty et al. 2019). Amino acid sequences are highly similar, with THCAS and CBCAS being 92% identical to each other and 84% and 83% identical to CBDAS, respectively (Sirikantaramas et al. 2004; Taura, Sirikantaramas, Shoyama, Yoshikai, et al. 2007; Laverty et al. 2019).

THCAS, CBDAS and CBCAS are members of the berberine bridge enzyme (BBE)-like gene family (PF08031) (Sirikantaramas et al. 2004; Taura, Sirikantaramas, Shoyama, Yoshikai, et al. 2007). This family is named after an oxidocyclase from *Eschscholzia californica* involved in alkaloid biosynthesis and part of the larger oxygen-dependent FAD-linked oxidoreductase family (PF02913) (Hauschild et al. 1998; Winkler et al. 2008). Like other BBE-like synthases, THCAS, CBDAS, and CBCAS contain an N-terminal signal peptide, a flavin adenine dinucleotide (FAD)-binding domain, a substrate-binding domain, and a BBE-like specific C-terminus that is part of the FAD-binding module (Sirikantaramas et al. 2004; Taura, Sirikantaramas, Shoyama, Yoshikai, et al. 2007). In accordance with this domain structure, THCAS and CBDAS have been found to be catalytically active in the extracellular storage cavity of the glandular trichome and rely on covalently bound FAD and O_2_ for their activity (Sirikantaramas et al. 2005; Rodziewicz et al. 2019). CBCAS is less extensively studied, but considering its high sequence similarity with THCAS, probably shares these biochemical activities (Morimoto et al. 1997; Taura, Sirikantaramas, Shoyama, Shoyama, et al. 2007; Gülck and Møller 2020). However, the latest phylogenetic classification of plant BBE-like genes was based on Arabidopsis sequences only (Brassicaceae) (Daniel et al. 2016) and consequently lacks genes from *Cannabis* and related genera. Even though some BBE-like enzymes related to cannabinoid oxidocyclases have been identified (Aryal et al. 2019), it still remains unclear exactly how the various described cannabinoid oxidocyclase genes are related to each other and to other BBE-like enzymes. Therefore, a comprehensive phylogenetic analysis of BBE-like enzymes including cannabinoid oxidocyclase genes is warranted.

### Cannabinoid oxidocyclase gene evolution

Although environmental factors play a role in determining the amount of cannabinoids present in different parts and stages of the plant (Rustichelli et al. 1998), in most populations the ratio between THCA and CBDA has been found to be under genetic control (Mandolino et al. 2003; Weiblen et al. 2015; Toth et al. 2020; Wenger et al. 2020). Codominant inheritance of CBDA and THCA chemotypes is consistent with a Mendelian single-locus (de Meijer et al. 2003; Onofri et al. 2015; Weiblen et al. 2015). This led to the model in which THCAS and CBDAS are encoded by alternate alleles of the same gene (B_T_ and B_D_, respectively) (de Meijer et al. 2003). However, later genome sequencing revealed that they are encoded by different genes (rather than alleles) within a large polymorphic genomic region with low levels of recombination (Kojoma et al. 2006; van Bakel et al. 2011; McKernan et al. 2015; Onofri et al. 2015; Weiblen et al. 2015; Grassa et al. 2018; Laverty et al. 2019). Thus, they are treated as separate genes below.

The genes encoding THCAS, CBDAS, and CBCAS have been identified (Sirikantaramas et al. 2004; Taura, Sirikantaramas, Shoyama, Yoshikai, et al. 2007; Laverty et al. 2019). The *THCAS* gene comprises a 1638 bp intronless open reading frame that is found in all drug-type cultivars (Sirikantaramas et al. 2004; Kojoma et al. 2006; van Bakel et al. 2011; McKernan et al. 2015; Onofri et al. 2015; Weiblen et al. 2015; Vergara et al. 2019). For this reason, the gene has been used as a diagnostic marker for detecting psychoactive cultivars for crop breeding and forensics (Kojoma et al. 2006; Kitamura et al. 2016). It should be noted, however, that a non-psychoactive cultivar from Malawi has a *THCAS* gene but accumulates the cannabinoid precursor CBGA instead of THCA. This is probably due to a single amino acid mutation leading to a defective (B_T0_) variant (Onofri et al. 2015). Gene copy number variation has been suggested based on amplicon sequencing of the *THCAS* locus (McKernan et al. 2015; Weiblen et al. 2015; Vergara et al. 2019). But amplicons may have included closely related genes such as *CBCAS* (see below). Thus, it remains unclear if *THCAS* occurs in multiple copies and, consequently, if copy number variation could be a target for breeding of cultivars.

The *CBDAS* gene comprises a 1632 bp intronless open reading frame that is found in all CBDA-producing cultivars (Taura, Sirikantaramas, Shoyama, Yoshikai, et al. 2007). However, different missense mutations have been described from CBGA-dominant hybrid cultivars from Italy and Ukraine that are considered B_D01_ and B_D02_ variants, respectively (Onofri et al. 2015). Another missense mutation was described from a cultivar from Turkey that is considered a weak B_Dw_ variant resulting in a partial accumulation of CBGA due to partly impaired activity of the encoded CBDAS (Onofri et al. 2015). In THCA-dominant cultivars, fragments have been found that are 93% identical to functional *CBDAS* and share a four base pair deletion that results in a truncated and most probably nonfunctional protein (Weiblen et al. 2015; Cascini et al. 2019; Vergara et al. 2019). This deletion showed strict association with THCA-producing cultivars, suggesting tight genetic linkage with *THCAS*. Indeed, it could be used to discriminate between fiber and drug-type cultivars as well as accurately predict chemotype in feral and cultivated plants (Cascini et al. 2019; Wenger et al. 2020). Notably, up to three different variants of such putative pseudogenes were detected in single cultivars (van Bakel et al. 2011; Weiblen et al. 2015; Laverty et al. 2019; Vergara et al. 2019) suggesting multiple duplicated loci. Moreover, given their low level of identity with *CBDAS* and close linkage with *THCAS* it is unclear if they should be considered variants of *CBDAS* or as separate loci.

The *CBCAS* gene comprises a 1638 bp intronless open reading frame. It was recently identified and described based on genome sequencing of the cultivar ‘Finola’ (Laverty et al. 2019). Based on sequence comparisons, “inactive THCA” synthase sequences described from European fiber-type cultivars more than a decade earlier were also considered CBCAS (Kojoma et al. 2006; Laverty et al. 2019). Other reported amplified fragments may also represent the same gene. For example, fragments from CBDA-dominant cultivars such ‘Carmen’ and ‘Canna Tsu’ had been reported to be similar to those “inactive THCA” synthases (McKernan et al. 2015; Weiblen et al. 2015). Similar lowly expressed fragments were also reported from other CBD-dominant cultivars and speculated to correspond with the “inactive THCA” synthases (Onofri et al. 2015). In order to confirm whether these indeed represent *CBCAS*, an meta-analysis is required.

In addition to *THCAS, CBDAS*, and *CBCAS*, other yet uncharacterized sequences have been described. These may encode enzymes for other cannabinoids, but their copy numbers and sequence properties are not well described or cataloged (Hurgobin et al. 2020). Besides the functionally characterized *CBDAS* gene, Taura *et al*. amplified two other gene fragments both of which contain a 1635 bp intronless open reading frame from genomic DNA and named these CBDAS2 and CBDAS3. They share 84% identity with *CBDAS* but did not encode enzymes with CBDAS activity (Taura, Sirikantaramas, Shoyama, Yoshikai, et al. 2007). Weiblen and coworkers amplified a fragment from cultivar ‘Carmen’ with 95% identity to *THCAS* and two identical fragments from cultivars ‘Skunk#1’ and ‘Carmen’ with 92% identity to *THCAS* (Weiblen et al. 2015). A pseudogene was described from the genome of cultivar ‘Purple Kush’ with 92% identity to *THCAS* but appeared phylogenetically separate from the previously mentioned fragments (van Bakel et al. 2011; Weiblen et al. 2015). In cultivar ‘Finola’, a putative pseudogene with 93% identity to *THCAS* was reported (Laverty et al. 2019). In various other fiber-type cultivars, “mutated *THCAS*” fragments were reported, some of which were pseudogenized (Cascini et al. 2019). A recent phylogenetic analysis also identified a set of lineages representing functional and nonfunctional “unknown cannabinoid synthases” (Vergara et al. 2019). But it remains unclear how these relate to the gene fragments listed above and to each other.

The total number of cannabinoid oxidocyclase genes varies considerably across cultivars. Onofri *et al*. amplified up to 5 (in cultivar ‘Haze’) different full-length fragments in drug-type cultivars and up to 3 (in cultivars from Yunnan and Northern Russia and an inbred Afghan hashish landrace) different full-length fragments in fiber-type cultivars (Onofri et al. 2015). Inbred individuals of cultivars ‘Carmen’ and ‘Skunk #1’ are expected to be homozygous but yielded four and five cannabinoid synthase fragments, respectively (Weiblen et al. 2015). McKernan *et al*. detected up to six different fragments (including pseudogenes) of *THCAS* and related sequences (McKernan et al. 2015). A recent study on copy number variation in cannabinoid oxidocyclase genes estimated that some of the analysed cultivars could have up to 10 different fragments (Vergara et al. 2019). Based on these results it is clear that cannabinoid oxidocyclase genes can be considered a unique gene family that stems from a recent expansion and includes genes with unknown function (Onofri et al. 2015; Weiblen et al. 2015; Vergara et al. 2019; Hurgobin et al. 2020). However, due to differences in i) primers used for amplification, ii) reference genomes used for copy number estimation, and iii) level of homozygosity, these numbers are not directly comparable and may not be accurate assessments of gene copy number. There is also no appropriate classification to facilitate the unequivocal naming and referencing of cannabinoid oxidocyclase genes.

Finally, it remains unclear whether these genes are specific to *Cannabis*. A phylogenetic analysis sampling cannabinoid oxidocyclase genes from cultivars ‘Skunk#1’, ‘Carmen’, and ‘Purple Kush’ *a priori* considered cannabinoid oxidocyclase genes to comprise a clade (Weiblen et al. 2015). A more recent phylogenetic analysis based on genomic data applied the same a priori assumption (Hurgobin et al. 2020). Another study sampling cultivars ‘Pineapple Banana Bubble Kush’ and ‘Purple Kush’ suggested that all cannabinoid oxidocyclase genes may comprise a clade but did not include functional *CBDAS* sequences nor homologs from *Cannabis’* most closely related genus *Humulus* (Vergara et al. 2019). Therefore, it remains unknown whether cannabinoid oxidocyclase genes are specific to *Cannabis* or represent more ancient gene duplications in e.g. an ancestor of *Cannabis* and related genera within the Cannabaceae family such as *Humulus* and *Trema (Padgitt-Cobb et al. 2019; Vergara et al. 2019)*. In addition, the genomic localization of many described gene sequences remains unknown and, consequently, we lack a clear overview of the patterns of gene duplication and divergence across the *Cannabis* genome (Weiblen et al. 2015).

### Aims

We present a comparative analysis of cannabinoid oxidocyclase genes in the genomes of *Cannabis*, closely related genera and informative outgroup species. This was greatly aided by the recent release and publication of several diverse *Cannabis* genome assemblies based on long-read sequencing technologies (Grassa et al. 2018; McKernan et al. 2018; Laverty et al. 2019; S. Gao et al. 2020). In addition, genomic information is available for other genera in the Cannabaceae family. Recent species-level phylogenetic analyses of the Cannabaceae family based on plastome sequences suggest that the genera *Parasponia* and *Trema* together are sister to *Cannabis* and *Humulus* (Jin et al. 2019). Draft genome assemblies have recently become available for *Humulus, Parasponia*, and *Trema*, that can be used for comparative analyses of *Cannabis* genes (van Velzen et al. 2018; Padgitt-Cobb et al. 2019; Kovalchuk et al. 2020). This provides an excellent opportunity to perform a comprehensive reconstruction of the evolution of cannabinoid oxidocyclase genes. In addition, *Morus notabilis* (Moraceae), *Medicago truncatula* (Fabaceae), and *Arabidopsis thaliana* (Brassicaceae) were included as outgroups, allowing us to place our results within a broader phylogenetic perspective and in the context of the existing BBE-like gene family classification (Daniel et al. 2016). Based on phylogenetic and synteny analysis, we elucidate the evolution of these genes and address the following questions:

1. How are cannabinoid oxidocyclases related to other berberine bridge enzymes?
2. Are cannabinoid oxidocyclase genes specific to *Cannabis* or do they represent more ancient duplications in e.g. an ancestor of *Cannabis* and related genera within the Cannabaceae family?
3. What are the phylogenetic relationships of *THCAS, CBDAS*, and *CBCAS* with other closely related genes?
4. What are the patterns of duplication and divergence of cannabinoid oxidocyclase genes across *Cannabis* genomes?

We also present a comprehensive clade-based classification of all cannabinoid oxidocyclase genes to resolve current confusion due to inconsistencies in naming and aid their future referencing and identification.

## Results

### Cannabinoid oxidocyclase genes are specific for *Cannabis*

To place cannabinoid oxidocyclase genes within the context of the BBE-like gene family we performed a phylogenetic analysis of BBE-like protein sequences from selected Eurosid genomes. These include genomes from *Cannabis sativa* cultivar ‘CBDRx’, *Humulus japonicus* cultivar ‘Cascade’, *Parasponia andersonii*, and *Trema orientale* from the Cannabacae family. Genomes from *Morus notabilis* (Moraceae), *Medicago truncatula* (Fabaceae), and *Arabidopsis thaliana* (Brassicaceae) were included as outgroups (Figure 2A). The resulting gene tree recovered eleven clades, including groups 1-7 earlier described (Daniel et al. 2016) based on Brassicaceae sequences (all seven groups were monophyletic, except that group 3 was confirmed to be nested within group 4). *Cannabis* BBE-like sequences were found in groups 2, 5.1, 5.2, 6, and 7.1. In addition, *Cannabis* sequence accession XP_030480925.1 represented an undescribed clade which we named group 8; *Cannabis* sequence accession XP_030480615.1 represented an undescribed clade including berberine bridge enzyme originally described from *Eschscholzia californica* which we named group 9. *THCAS, CBDAS*, and *CBCAS* are members of a newly defined group 10. Within this group, a Cannabaceae-specific gene expansion can be identified within which all three known cannabinoid oxidocyclase occur in a *Cannabis*-specific clade, which we name the cannabinoid oxidocyclase clade. This suggests that THCAS, CBDAS, and CBCAS originated from a single ancestral cannabinoid oxidocyclase gene within the *Cannabis* lineage.

**Fig 2.**
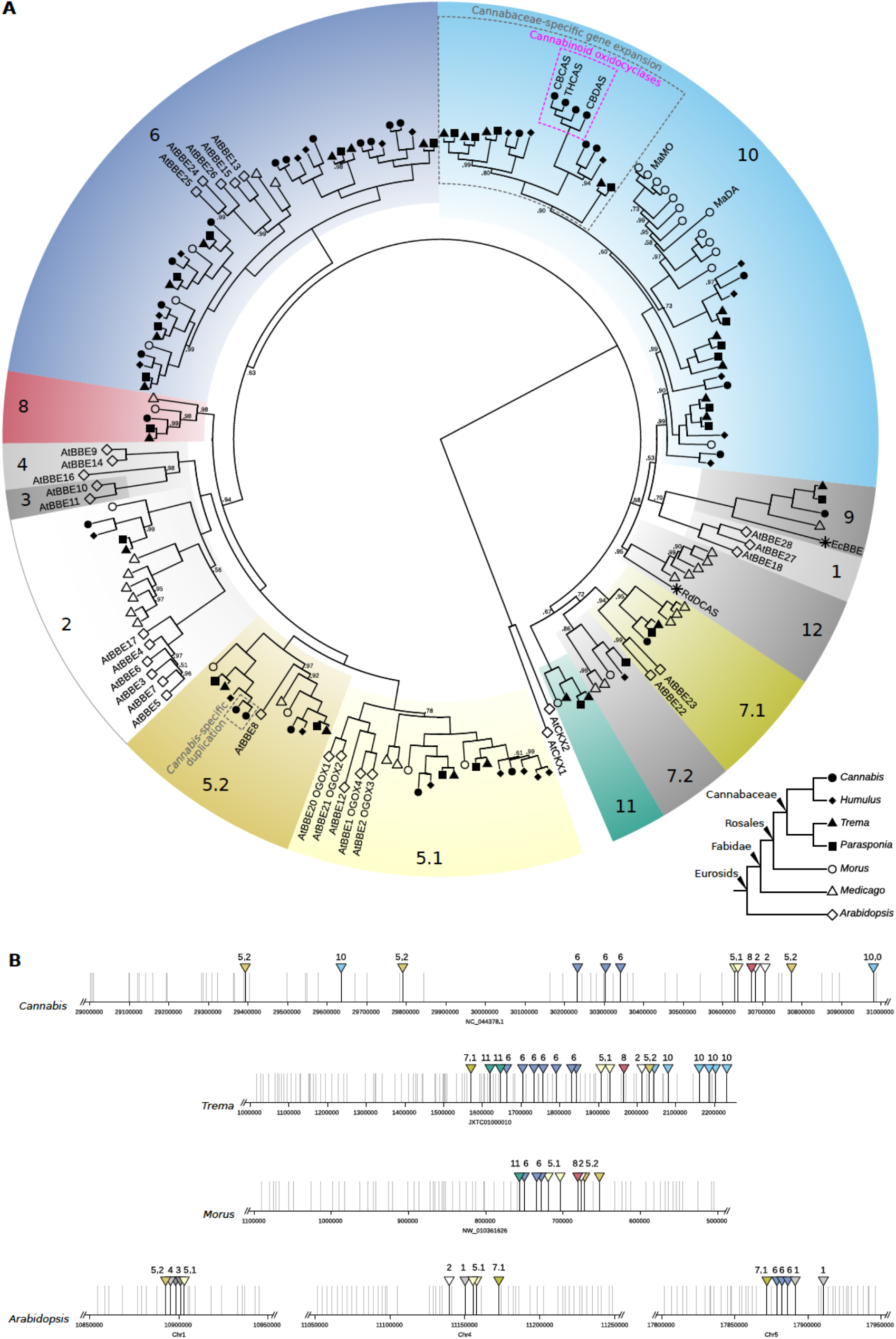
Eurosid berberine bridge enzyme gene family analysis. A. Gene tree based on protein sequences showing that cannabinoid oxidocyclases comprise a *Cannabis*-specific clade. Shapes indicate sampled species *Cannabis sativa* (solid circles), *Humulus lupulus* (solid diamonds), *Parasponia andersonii* (solid squares), *Trema orientale* (solid triangles), *Morus notabilis* (open circles), *Medicago truncatula* (open triangles), and *Arabidopsis thaliana* (open diamonds), stars indicate additional sequences from *Rhododendron dauricum* and *Eschscholzia californica*. Coloured blocks indicate the identified groups 1-12; node labels indicate posterior probabilities below 1.0. Bottom right inset shows known relationships among sampled species. B. genomic colocalization of berberine bridge enzymes in *Cannabis sativa* cultivar ‘CBDRx’, *Trema orientale, Morus notabilis*, and *Arabidopsis thaliana*. Grey horizontal bars indicate contigs in the *Cannabis* chromosomal scaffold shown in Figure 4A. Vertical lines indicate locations of annotated genes; berberine bridge enzymes are indicated with triangles in colour consistent with those in panel A. For displaying purposes, genomic scaffolds are not shown in the same scale (size shown is indicated).

We also found that BBE-like genes often occurred near each other in the *Cannabis* CBDRx genome. We therefore retrieved genomic locations of all BBE-like genes in other genomes including *T. orientale, M. notabilis*, and *A. thaliana*. This revealed that BBE-like genes from different clades are commonly colocalized in these genomes (Figure 2B). This suggests that selection favors BBE-like genes to remain in close genomic proximity. It is known that genes involved in the same pathway have the tendency to cluster in plant genomes (Liu et al. 2020). However, it is not clear if and how the various BBE-like genes share pathways and we therefore have no conclusive explanation for this intriguing pattern.

### Phylogenetic classification of cannabinoid oxidocyclase genes

To elucidate the phylogenetic relationships of THCAS, CBDAS, and CBCAS with other highly homologous genes, we performed an extensive phylogenetic analysis of cannabinoid oxidocyclase genes from genome assemblies of *Cannabis* cultivars ‘CBDRx’, ‘Jamaican Lion’, ‘Finola’, ‘Purple Kush’, and a putatively wild *Cannabis* plant from Jilong, Tibet (Grassa et al. 2018; McKernan et al. 2018; Laverty et al. 2019; S. Gao et al. 2020; McKernan et al. 2020). Additional analyses based on sequences from NCBI and from more fragmented *Cannabis* genomes are shown in Figures S1 and S2, respectively. Based on the resulting gene trees we consistently recovered the same three main clades (A-C; Figures 3) that we describe below.

**Fig 3.**
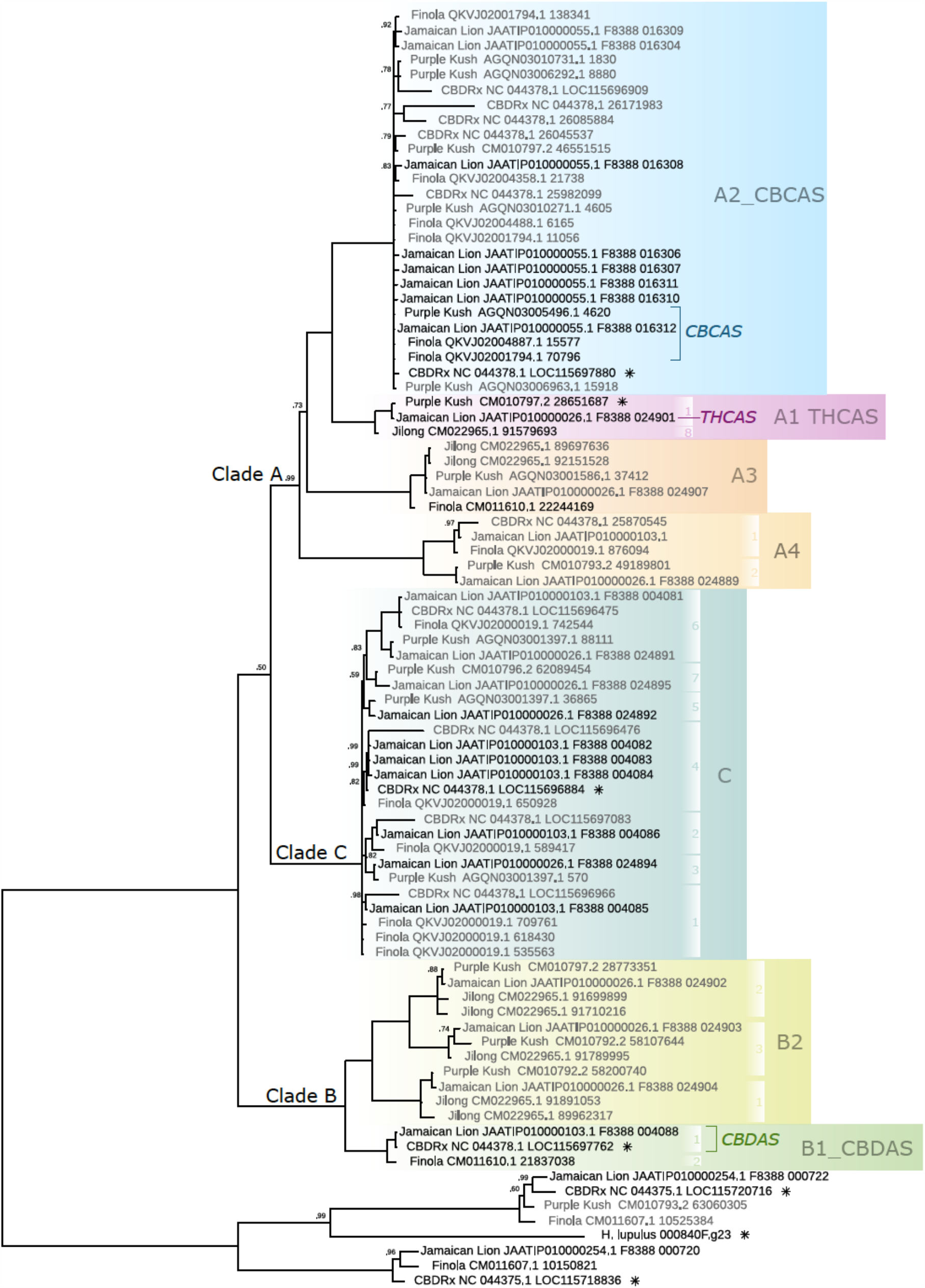
Cannabinoid oxidocyclase gene tree. Based on nucleotide sequences from whole-genome assemblies of *Cannabis sativa* cultivars ‘CBDRx’, ‘Finola’, ‘Jamaican Lion’, and a wild plant from Jilong, Tibet (Grassa et al. 2018; McKernan et al. 2018; Laverty et al. 2019; S. Gao et al. 2020). Labels indicate genbank accession of genomic contig and locus tag (when available) or start position. Putative nonfunctional (pseudo)genes are in grey; functionally characterized THCAS, CBDAS, and CBCAS genes (Sirikantaramas et al. 2004; Taura, Sirikantaramas, Shoyama, Yoshikai, et al. 2007; Laverty et al. 2019) are labeled. Sequences included in the BBE-like analysis shown in Figure 2A are marked with an asterisk. Coloured blocks indicate the identified clades; white blocks indicate sequence types. Node labels indicate posterior probabilities below 1.0.

#### Clade A

Clade A includes *THCAS* and *CBCAS*. It comprises four subclades and can be characterized by three unique nonsynonymous substitutions (Table S1). Subclade A1_THCAS comprises full-length coding sequences from THCA-producing plants such ‘Purple Kush’, ‘Skunk#1’, and ‘Chemdog91’, including functionally characterised THCAS (Sirikantaramas et al. 2004), ‘‘drug-type’’ THCAS sequences (Kojoma et al. 2006), “active’’ THCAS sequences (McKernan et al. 2015), and fully functional (B_T_) as well as nearly defective (B_T0_) coding sequences (Onofri et al. 2015) (Figures 3, S1, S2). Subclade A1_THCAS sequences can be characterized by three unique nonsynonymous substitutions and may be further divided into 2 groups and 18 types of which 9 were previously described by (Onofri et al. 2015) and 9 are new (Table S1). Group 1 comprises six types (1/1-1/6) that are identical or similar to the *THCAS* reference (Sirikantaramas et al. 2004; Onofri et al. 2015). Type 1/1 (Onofri et al. 2015) comprises sequences from various cultivars that are identical to the functionally characterized THCAS described by (Sirikantaramas et al. 2004). Type 1/2 (Onofri et al. 2015) comprises accession KP970849.1 which differs by a single synonymous substitution from type 1/1 and can therefore be considered functionally equivalent. Type 1/3 (Onofri et al. 2015) comprises the defective B_T0_ allele from Malawi that differs only by 706C (Gln). Type 1/4 (Onofri et al. 2015) comprises sequences from cultivars ‘Skunk #1’, ‘AK47’, ‘Chemdog91’, ‘CannaTsu’, ‘Black84’, and a hashish landrace from Afghanistan that share 749A (Asp). Type 1/5 comprises sequences from cultivars ‘Purple Kush’, ‘Blueberry Essence’, and ‘C4xCannaTsu’ that share the unique amino acid (aa) substitution 998G (Arg). Type 1/6 comprises accession KT876046.1 from cultivar ‘Otto’ that differs by only one aa substitution. Group 2 comprises six types sharing the nonsynonymous substitution 373C (Leu). Type 2/1 (Onofri et al. 2015) comprises sequences from cultivars ‘Haze’, ‘Alaskan ice’, and ‘Otto’ that share two nonsynonymous substitutions. Type 2/2 (Onofri et al. 2015) comprises accession KP970853.1 from cultivar ‘Haze’ that has one nonsynonymous substitution. Type 2/3 (Onofri et al. 2015) comprises accession which differs by two synonymous substitutions from type 2/1 and can therefore be considered functionally equivalent. Type 2/4 comprises sequences from Boseung province in Korea described by (Doh et al. 2019) that share three nonsynonymous substitutions. Type 2/5 comprises accession MN422091.1 from Jecheon province in Korea that has five nonsynonymous substitutions. Type 2/6 comprises sequences from low-THCA cultivar ‘Cheungsam’ described by (Doh et al. 2019) and can be characterised by ten nonsynonymous substitutions. Other types remain ungrouped. Type 3 (Onofri et al. 2015) comprises accession KP970851.1 from a hashish landrace from Afghanistan that has the unique aa substitution 187C (Leu). Type 4 (Onofri et al. 2015) comprises accession KP970855.1 from cultivar ‘Haze’ and has aa substitutions 794G (Gly) and 1229A (Glu). Type 5 comprises partial sequences from various regions in Pakistan described by (Ali et al. 2019) of which at least one is a pseudogene. They can be characterized by two unique aa substitutions: 851T (Val) and 883C (Pro). Type 6 comprises accession MT338560.1 from Oregon CBD that shares 998G (Arg) with type 1/5 but has an additional unique aa substitution 1064A (Asn). Type 7 comprises accession LC120319.1 from cultivar ‘Big Bud’ described by (Kitamura et al. 2016) that shares 749A (Asp) with type 1/4 but has one additional nonsynonymous substitution (1018G; Ala). Type 8 comprises the putative *THCAS* sequence of a putatively wild plant from Jilong, Tibet that can be characterized by six unique aa substitutions.

Subclade A2_CBCAS comprises full-length coding as well as nonfunctional (pseudo)gene sequences from drug-type cultivars such as ‘Purple Kush’ and ‘Jamaican Lion’ as well as fiber-type cultivars such as ‘Finola’ and ‘Carmen’. It includes the functionally characterised *CBCAS* (Laverty et al. 2019), “mutated *THCAS”* (Cascini *et al*. 2019), ‘‘fiber-type *THCAS”* sequences (Kojoma *et al. 2006)*, and “inactive THCAS” sequences ((Kojoma et al. 2006; McKernan et al. 2015; Cascini et al. 2019; McKernan et al. 2020). Subclade A2_CBCAS sequences can be characterized by 12 unique aa substitutions (Table S1).

Subclade A3 comprises at least one full-length coding sequence from cultivar ‘Finola’ and at least two nonfunctional (pseudo)gene copies from drug-type cultivars ‘Purple Kush’ and ‘Jamaican Lion’. These sequences can be characterized by a duplication of the 3rd codon (TAC; Tyr) and 13 unique nonsynonymous substitutions (Table S1). They have not yet been functionally assessed but given that at least one variant comprises a full-length coding sequence it is expected to have some functional relevance; probably as a cannabinoid oxidocyclase.

Subclade A4 comprises three nonfunctional (pseudo)gene sequences from whole-genome assemblies of cultivars ‘Purple Kush’, ‘Finola’, and ‘Jamaican Lion’. They share four nonsense mutations and 14 unique aa substitutions (Table S1).

#### Clade B

Clade B comprises two subclades and can be characterized by 16 unique aa substitutions (Table S1). Subclade B1_CBDAS comprises full-length coding *CBDAS* sequences from CBDA-producing cultivars such as ‘Finola’, ‘Carmen’, and ‘CBDRx’ (Figures 3, S1, S2). These sequences can be characterized by a 3bp deletion at position 755 and 14 unique aa substitutions (Table S1). They can be further divided into two types that correspond with groups 5 and 6 described by (Onofri et al. 2015). Type 1 is characterized by 1423A (Lys) and found in fiber-type cultivars such as ‘Ermo’ and ‘C.S.’, drug-type cultivar ‘Jamaican Lion’, and landraces from China and Japan (Taura, Sirikantaramas, Shoyama, Yoshikai, et al. 2007; Onofri et al. 2015; Grassa et al. 2018; Cascini et al. 2019). It includes the CBDA synthase gene CBDAS1 and a defective sequence B_D01_ coding sequence (Taura, Sirikantaramas, Shoyama, Yoshikai, et al. 2007; Onofri et al. 2015). Type 2 can be characterized by three unique Serines (Table S1) and is found in fiber-type cultivars such as ‘Finola’, ‘Carmen’, ‘Ermes’, ‘Futura 75’, ‘Tygra’, and ‘Uso31’ (Onofri et al. 2015; Weiblen et al. 2015; Cascini et al. 2019; Laverty et al. 2019). It includes fully functional (B_D_), weakly functional (B_DW_) and defective (B_D01_ and B_D02_) coding sequences (Onofri et al. 2015). We note that some sequences described by Cascini *et al*. are ambiguous at type-specific positions and so probably represent a mix of both types (see Table S3).

Subclade B2 comprises the nonfunctional (pseudo)gene sequences from THCA-producing cultivars such as ‘Purple Kush’, ‘Jamaican Lion’, ‘Skunk#1’, ‘Chocolope’, and ‘Northern Light’; including “mutated CBDAS” sequences described by Cascini *et al*. and “marijuana-type CBDA synthase” sequences described by Weiblen et al. (Weiblen et al. 2015; Cascini et al. 2019) (Figures 3, S1, S2). They can be characterized by a 4 or 6 bp frame-shift deletion at position 153 and by two unique nonsynonymous substitutions (Table S1). They can be further divided into three types based on additional missense and aa mutations (we note, however, that because of the shared frame-shift deletions these mutations probably did not have any significance in terms of actual coding sequence and can therefore be considered “secondary”). The first type can be characterised by four secondary nonsense mutations and eight secondary unique aa substitutions (Table S1). The second type can be characterised by two secondary unique aa substitutions. The third type can be characterised by nine secondary unique aa substitutions (Table S1). Accession LKUA01006620.1 from cultivar ‘LA confidential’ may be a chimera of types 2 and 3.

#### Clade C

Clade C comprises full-length coding as well as nonfunctional (pseudo)gene sequences from cultivars ‘Purple Kush’, ‘Finola’, ‘CBDRx’, and ‘Jamaican Lion’ as well as *CBDAS2* and *CBDAS3* from a ‘domestic’ cultivar from Japan described as having no CBDA synthase activity by Taura *et al*. (Taura, Sirikantaramas, Shoyama, Yoshikai, et al. 2007) (Figures 3, S1, S2). They share 19 unique aa substitutions and can be divided into 7 types (Figure 3; Table S1).

### Patterns of cannabinoid oxidocyclase gene duplication and divergence

To reconstruct patterns of gene duplication and divergence, we assessed microsynteny across genomes of *Cannabis* cultivars ‘CBDRx’, ‘Jamaican Lion’, ‘Finola’, ‘Purple Kush’, and a putatively wild *Cannabis* plant from Jilong, Tibet. Based on nucleotide alignments and protein comparisons, we found that all cannabinoid oxidocyclase genes occur in two main syntenic clusters, together with other BBE-like genes. The first main syntenic cluster comprises a tandemly repeated array of genes from clade C in the genome assemblies of cultivars ‘Finola’, ‘CBDRx’, and ‘Jamaican Lion’ (Figure 4A). The array is flanked at the 3’ end by a group 5.2 BBE-like gene, a receptor-like protein, a Patellin protein, a TWINKLE DNA primase-helicase, and a caseinolytic protease. In the Jamaican Lion genome, there are two putative allelic variants; first comprising two full-length coding sequences and two nonfunctional (pseudo)gene copies and the second comprising five full-length coding sequences and one nonfunctional (pseudo)gene copy. In the assembly of cultivar ‘Finola’, it comprises six nonfunctional (pseudo)gene copies. In the assembly of cultivar ‘CBDRx’, it comprises one full-length coding sequence and four nonfunctional (pseudo)gene copies. The array is flanked at the 5’ end by one of three variants of a large genomic region with very little nucleotide-level identity (Figure S5). All variants comprise another copy of a group 5.2 BBE-like gene. The first variant comprises a single copy of *THCAS*, a tandemly repeated array of subclade B2 nonfunctional (pseudo)genes, and a nonfunctional (pseudo)gene from the A4 subclade. It is present in cultivars ‘Jamaican Lion’, ‘Purple Kush’, and the putatively wild plant from Jilong, Tibet. The second variant comprises a single copy of a type 1 *CBDAS* and can be found in cultivars ‘Jamaican Lion’, and ‘CBDRx’. The third variant comprises only a type 2 *CBDAS* and can be found in cultivar ‘Finola’. We found no nucleotide-level alignments between these variants except for the context around the group 5.2 BBE-like in the second and third variants (Figures 4A and S5). This suggests high levels of divergence across this large genomic region.

**Fig 4.**
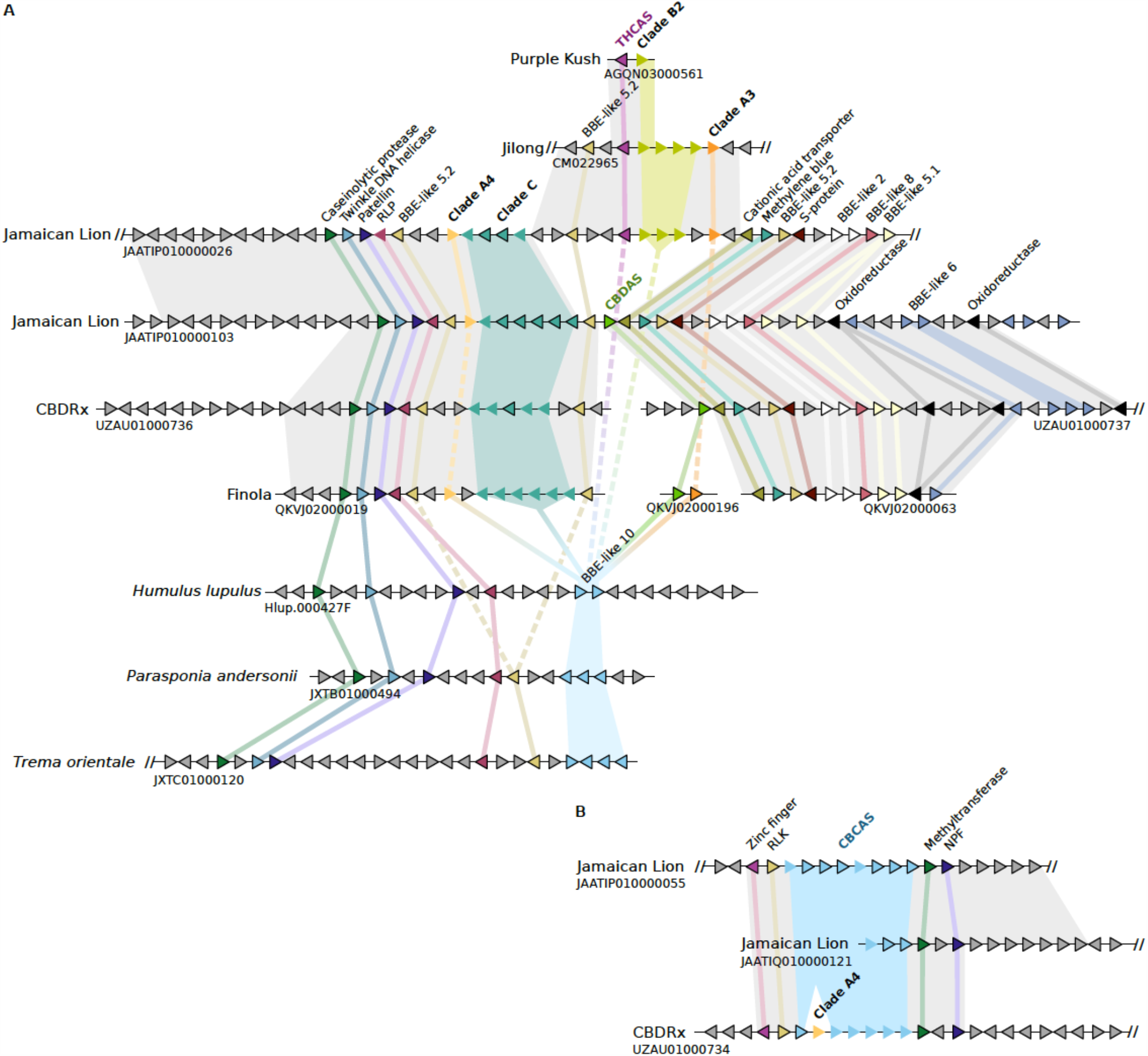
Cannabinoid oxidocyclase microsynteny assessments. A. syntenic block comprising CBDAS, THCAS, and related genes. B. syntenic block comprising CBCAS tandemly repeated array. Triangles indicate genes (not to scale) coloured according to their homology and putative orthologs are connected with coloured lines. Nonfunctional (pseudo)genes are shown without black outlines. Grey backgrounds indicate LASTZ nucleotide alignments based on results shown in Figs. S3-4. Cannabinoid oxidocyclase genes are members of BBE-like group 10 (see Fig. 2) and labeled in boldface. Abbreviations: BBE, berberine bridge enzyme; NPF, NRT1 / PTR family protein; RLK, receptor-like kinase; RLP, receptor-like protein.

The second cluster comprises a tandemly repeated array of genes from subclade A2_CBCAS in the genome assemblies of cultivars ‘CBDRx’ and ‘Jamaican Lion’ (Figure 4B). The array is flanked at the 5’ end by a RING/FYVE/PHD-type zinc finger family protein and a receptor-like kinase; and at the 3’ end by an ankyrin repeat family protein and an NRT1 / PTR family protein in both assemblies. Some of these flanking genes are considered pathogen response genes (McKernan et al. 2020). In the ‘Jamaican Lion’ genome, there are two putative allelic variants, the first of which comprises six full-length *CBCAS* coding sequences and two nonfunctional (pseudo)gene copies and the second of which is only partially assembled. In the ‘CBDRx’ genome, it comprises one full-length *CBCAS* coding sequence and five nonfunctional (pseudo)gene copies. Interestingly, it also includes a nonfunctional (pseudo)gene from subclade A3 but given the lack of additional A3 copies within the array, this appears to be a relatively recent insertion. In the genome assemblies of cultivars ‘Finola’ and ‘Purple Kush’, subclade A2_CBCAS gene copies appear in several unplaced scaffolds probably representing the same array (Hurgobin et al. 2020). Subclade A2_CBCAS gene copies are absent in the assembly of the putatively wild plant from Tibet. No further synteny was found with *Humulus, Parasponia*, or *Trema*; suggesting that this syntenic cluster is specific to *Cannabis*.

To assess the direction of evolution, we then assessed protein-level microsynteny in genomes from closely related Cannabaceae species *H. lupulus, P. andersonii*, and *T. orientalis*. This revealed that each comprises a tandemly repeated array of group 10 BBE-like genes that are closely related to the known cannabinoid oxidocyclase genes, as well as a single copy of the group 5.2 BBE-like gene (not found in *Humulus*) and the receptor-like protein, the Patellin protein, the TWINKLE DNA primase-helicase, and the caseinolytic protease listed above (Figure 3A). This suggests that cannabinoid oxidocyclases originated within an ancestral syntenic block and experienced a series of tandem gene duplications, translocations, and divergence.

## Discussion

### Origin of cannabinoid oxidocyclases from within the BBE-like gene family

Since the cannabinoid oxidocyclase genes were first discovered and described, it has been known that they are members of the BBE-like gene family (Sirikantaramas et al. 2004; Daniel et al. 2017). However, the BBE-like family is large and the most recent classification of plant BBE-like genes was based only on analysis of genes from *Arabidopsis* in the Brassicaceae family (Daniel et al. 2016). Even though some BBE-like enzymes related to cannabinoid oxidocyclases have been identified (Aryal et al. 2019), it remained unclear how the various described cannabinoid oxidocyclase genes are related to each other and to other berberine bridge enzymes.

Our results show that cannabinoid oxidocyclase genes from *Cannabis* originated from a newly defined clade (Group 10) within the BBE-like gene family (Figure 2A). *Rhododendron dauricum* daurichromenic acid synthase (*RdDCAS*), another plant cannabinoid oxidocyclase (Iijima et al. 2017), originates from another clade (Group 12). Within Group 10 gene expansions occurred independently in Moraceae and Cannabaceae (Figure 2A). The expansion in Moraceae includes the recently described *Morus alba* Diels–Alderase (*MaDA*) and moracin C oxidase (*MaMO*) genes that are responsible for the production of the medicinal compound chalcomoracin (Han et al. 2018; L. Gao et al. 2020). The expansion in Cannabaceae eventually led to the origin of cannabinoid oxidocyclases. Such gene diversification and enzymatic versatility confirms that BBE-like enzymes play important roles in generating biochemical novelty (Daniel et al. 2017). Because most plant BBE-like genes (including those in Group 10) have a secretory signal peptide they may be considered to be ‘pre-adapted’ for a role in the extracellular space. Within the Cannabaceae-specific gene expansion, *THCAS, CBDAS*, and *CBCAS* form a clade that is sister to homologous genes from *Cannabis* and *Humulus* (Figure 2A). These results show unequivocally and for the first time that cannabinoid oxidocyclase genes did not originate from more ancient duplications within the Cannabaceae but are specific to *Cannabis*.

The central cannabinoid precursor CBGA is the common substrate for THCAS, CBDAS, and CBCAS (Figure 1). We therefore hypothesise that the CBGA biosynthetic pathway existed before the origin of cannabinoid oxidocyclases. Thus, other *Cannabis* cannabinoid biosynthesis genes such as those encoding CBGAS, OAC, OLS, and acyl-activating enzyme 1 (Raharjo et al. 2004; Taura et al. 2009; Gagne et al. 2012; Stout et al. 2012) may have originated from more ancient duplications in an ancestor of *Cannabis* and related genera within the Cannabaceae family such as *Humulus, Parasponia*, and *Trema*. The comparative approach that we leveraged here can help elucidate the order in which these pathway genes evolved and, thus, reconstruct the origin of a novel and societally relevant biosynthetic pathway.

### A novel classification of cannabinoid oxidocyclase genes

Based on our phylogenetic meta-analysis, we classified the cannabinoid oxidocyclase genes into three main clades (A-C) comprising a total of seven (sub)clades (Figure 3). *THCAS* and *CBCAS* are most closely related and occur in subclades A1 and A2, respectively, while *CBDAS* occurs in subclade B1. In addition to these three subclades representing functionally characterized cannabinoid oxidocyclase genes, we identified four previously unrecognized subclades. Based on current sampling, two of these clades contain only pseudogenes. Within subclade A4, two types can be recognized that share four nonsense mutations. Similarly, within subclade B2, three types can be recognized that share a frame-shift mutation (Table S1). Therefore, it seems that within each of these two subclades, the most-recent common ancestor was likely already nonfunctional. Contrastingly, clade C and subclade A3 each include full-length coding sequences that are most likely functional enzymes. Taura *et al*. (2007) prepared recombinant enzymes based on two clade C full-length sequences (accession numbers AB292683.1 and AB292683.1) and reported they did not exhibit CBDA synthase activity but did not show the underlying experimental data. The subclade A3 sequence from cultivar ‘Finola’ was reported as a pseudogene (Laverty et al. 2019) but based on our assessment of the genome sequence deposited on genbank it encodes a full-length protein. It has not yet been experimentally tested. Consequently, there is potential that the products of clade C and subclade A3-encoded enzymes are of biochemical and potential medical importance.

Our clade-based classification is intended to aid unequivocal referencing and identification of cannabinoid oxidocyclase genes. For example, based on our analyses we were able to confirm that sequences variously named “Fiber-type”, “inactive”, or “obscure” *THCAS* (McKernan et al. 2015; Onofri et al. 2015; Weiblen et al. 2015; McKernan et al. 2020) can be classified as variants of *CBCAS* (Laverty et al. 2019). Similarly, we reclassify sequences variously described as “marijuana-type” or “mutated” *CBDAS* (Weiblen et al. 2015; Cascini et al. 2019) as representing subclade B2. In retrospect, much of the confusion about gene identity stems from the general tendency to name sequences in accordance with the primers used for their amplification. For example, CBCAS-like and clade B2 pseudogenes were probably erroneously classified because they were generally amplified with primers that were considered specific for *THCAS* or *CBDAS*, respectively. Sequences representing clade C have been often neglected in amplicon-based studies because they did not amplify using *THCAS* or *CBDAS* primers (Onofri et al. 2015). Moreover, these sequences were variously considered either a variant of *CBDAS* or *THCAS*, leading to additional confusion (Taura, Sirikantaramas, Shoyama, Yoshikai, et al. 2007; Weiblen et al. 2015). We anticipate that our classification will help avoid such confusion about the identity and relationships of cannabinoid oxidocyclase genes in the future. Our comprehensive meta-analyses sampling all currently available sequences consistently recovered the same clades (see Figures 3, S1, S2), suggesting that this classification is robust. New cannabinoid oxidocyclase sequences can be associated with the corresponding clade by phylogenetic analysis or based on the clade-specific missense mutations listed in Table S1. Our sequence alignments and phylogenetic trees are available for analysis via TreeBase/DRYAD. In case sequences fall outside any of our described clades, new (sub)clades can be defined in accordance with our system.

### Localization and divergence of oxidocyclase genes in the *Cannabis* genome

The genetic basis underlying the ratio of THCA and CBDA is relatively well known. Genome sequencing of the drug-type cultivar ‘Purple Kush’ and fiber-type cultivar ‘Finola’ revealed that THCAS and CBDAS genes are located at different loci within a single large polymorphic genomic region with low levels of recombination (Laverty et al. 2019). However, the genomic locations of most other oxidocyclase genes has remained unknown. Consequently, a comprehensive overview of the patterns of gene duplication and divergence across the *Cannabis* genome has been lacking (Weiblen et al. 2015). Our assessment of microsynteny based on nucleotide alignments and protein comparisons revealed that cannabinoid oxidocyclase genes occur in two large syntenic blocks. The first block comprises a conserved region including a tandemly repeated array of clade C genes and a divergent region including either THCAS and subclade B2 pseudogenes; CBDAS and a subclade A3 gene; or only CBDAS (Figure 4A, S3). Close linkage of clade B2 pseudogenes with THCAS explains why they are considered markers for drug-type cultivars (Cascini et al. 2019). It also explains why a “CBDAS genotype assay” differentiating between subclades B1 and B2 can be used to accurately predict levels of THCA versus CBDA (Wenger et al. 2020). The second block comprises a conserved region including a tandem repeat of CBCAS-like genes (Figure 4B). This explains why, even though CBCAS-like genes have been considered to be associated with fiber-type cultivars and genomes of some drug-type cultivars indeed lack any CBCAS-like gene, genomes of other drug-type cultivars such as ‘Purple Kush’, ‘Skunk #1’, and ‘Pineapple Banana Bubble Kush’ do have full-length CBCAS genes. We have not found any nonfunctional THCAS gene closely linked to CBDAS as predicted by (Wenger et al. 2020). However, given the deep divergence between THCAS and CBDAS it seems unlikely that they comprise orthologous genes (see Figures 3 and 4). Similarly, it is yet unclear if CBDAS and the subclade B2 pseudogenes are orthologous or comprise different paralogous loci. Long-read whole genome sequencing of additional cultivars or wild plants may uncover haplotypes including *THCAS, CBDAS* and clade B2 pseudogenes and help elucidate these aspects.

### An evolutionary model for the origin and diversification of cannabinoid oxidocycloase genes

Our protein-level microsynteny analysis including genomes from closely related species *H. lupulus, P. andersonii*, and *T. orientalis* revealed an ancestral syntenic block including several Group 10 BBE-like genes (Figure 4A). Below we propose our most parsimonious evolutionary interpretation of cannabinoid oxidocyclase gene duplication and divergence (Figure 5). First, a group 10 BBE-like gene in an ancestral *Cannabis* neofunctionalized to use CBGA as substrate. Subsequent gene duplication and divergence lead to a set of ancestral genes representing the three main extant clades A, B, and C. Next, tandem duplication of this set together with the closely linked group 5.2 BBE-like gene resulted in two blocks. Block 1 retained ancestral genes representing clades A (diverging into an ancestor of the extant clade A4 pseudogenes) and C; clade B was apparently lost. Block 2 retained ancestral genes representing clades A (an ancestral clade A3 gene originated through duplication and divergence) and B; clade C was apparently lost. Support for this hypothetical tandem duplication event can be found in the BBE-like gene tree where the two corresponding group 5.2 BBE-like genes also comprise a *Cannabis*-specific duplication (Figure 2A). Finally, large-scale divergence of the second block led to the three variants of the divergent region described above. The tandemly repeated array of subclade A2_CBCAS genes most likely originated from duplication of an ancestor of *THCAS* and translocation to another chromosomal region.

**Fig 5.**
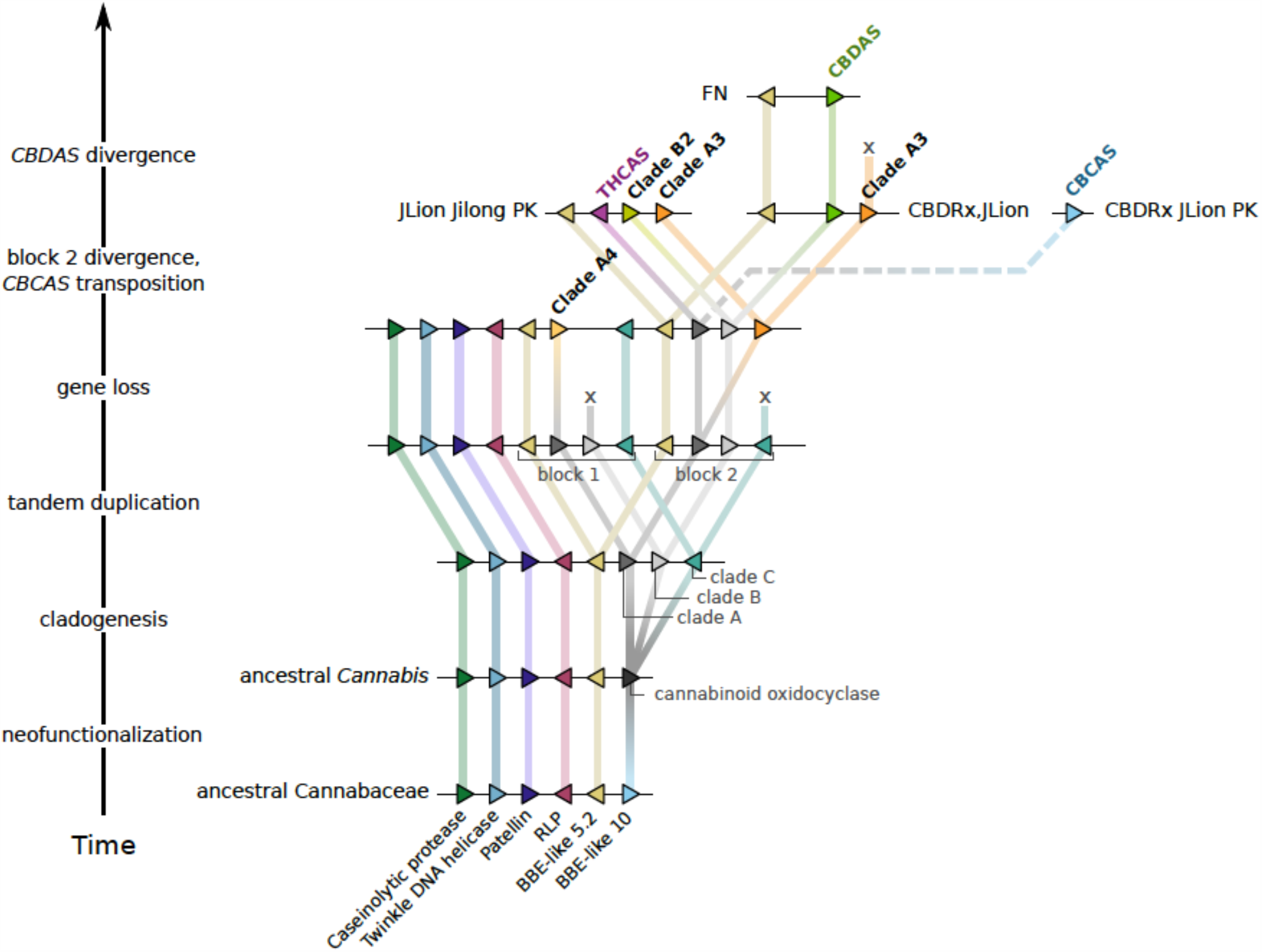
Evolutionary model of cannabinoid oxidocyclase gene duplication and diversification. Most parsimious hypothesis based on microsynteny patterns shown in Fig. 4 and phylogenetic reconstruction shown in Fig. 3. Triangles indicate genes (not to scale) coloured according to their homology and putative orthologs are connected with coloured lines. Dashed line indicates transposition of CBCAS to another syntenic block. Cannabinoid oxidocyclase genes are labeled in boldface. Abbreviations: BBE, berberine bridge enzyme; RLP, receptor-like protein.

The nature of the hypothesized ancestral cannabinoid oxidocyclase remains unknown (Vergara et al. 2019). Based on the observation that sequence variation was higher among “CBDAS”-like than among “THCAS”-like sequences, Onofri et al. considered CBDAS the ancestral type (Onofri et al. 2015). We now know, however, that this perceived variation was due to the existence of additional gene lineages (Figure 3). Based on our gene tree, it is not possible to conclusively reconstruct whether the ancestral function was similar to that of THCAS, CBDAS, CBCAS, or another yet unidentified synthase (Vergara et al. 2019). Nevertheless, given the relative recent divergence of cannabinoid oxidocyclase genes it may be possible to reconstruct an ancestral protein sequence for functional testing with reasonable accuracy.

### Re-evaluating gene copy number variation

Earlier studies have assessed cannabinoid oxidocyclase gene copy number with the aim to link this to chemical variation (McKernan et al. 2015; Weiblen et al. 2015; Vergara et al. 2019). We argue that, in light of our phylogenetic classification, these earlier results need to be reevaluated/reassessed. Claims have been made with regard to copy number variation in THCAS (McKernan et al. 2015; Weiblen et al. 2015; Vergara et al. 2019). However, we have found no instance of multiple copies of the subclade A1 functional THCAS gene. This is in line with recent findings based on a similar comparative genomics approach (Hurgobin et al. 2020). Instead, copies counted toward “THCAS” copy number variation were due to tandem repeat copies of subclade A2. Similarly, although multiple types can be recognized within subclade B1_CBDAS they have been recovered exclusively as a single copy. Instead, we found copy number variation reported for *CBDAS* to be due to amplification of tandem repeat copies of subclade B2 pseudogenes in THCA-producing cultivars (McKernan et al. 2015; Weiblen et al. 2015; Vergara et al. 2019). Tight genetic linkage of *THCAS* and subclade B2 tandemly repeated array (Figure 4) explains why “CBDAS copy number” was found to be positively correlated with the production of THCA and negatively with that of CBDA (Vergara et al. 2019).

Based on our classification, claims of *THCAS* and *CBDAS* copy number variation can be attributed to off-target amplification by the primers used (McKernan et al. 2015; Weiblen et al. 2015). Particularly illustrative in this regard is that sequences from clade C have been considered copies of *THCAS* or *CBDAS* in different studies, depending on the primers used for amplification (Taura, Sirikantaramas, Shoyama, Yoshikai, et al. 2007; Weiblen et al. 2015). Thus, based on currently available data, we consider *THCAS* and *CBDAS* each as single-copy genes and that gene copy number variation is restricted to the tandem duplications of subclades A2_CBCAS and B2, and clade C. Given that these (sub)clades may include a variable number of pseudogenes (Figure 4), it is not apparent how their copy number would have important functional relevance. For example, no correlation was found between copy number of CBCAS-like genes and the production of CBCA (Vergara et al. 2019). Another question is whether human selection has led to copy number variation in cannabinoid oxidocyclase genes (Weiblen et al. 2015; Vergara et al. 2019). Given the considerable divergence between the different subclades (Figure 3), we assume that their divergence most likely predated human selection. This is also true for the subclade B2 pseudogenes, which include three divergent types. Based on increased local dN/dS ratios in the ancestor of subclade B2, Weiblen *et. al*. suggested that human selection had favored these pseudogene to increase potency (Weiblen et al. 2015). We would rather attribute any such selection to the *THCAS* gene that is directly responsible for potency and to which the tandemly repeated array of subclade B2 genes is closely linked (Figure 4). But increased dN/dS could also simply be attributed to the fact that pseudogenes are generally considered to be evolving neutrally (Li et al. 1981). Tandem repeats of highly similar paralogs such as those within subclade A2 and clade C tandemly repeated arrays could have originated recently (Figures S3, S4). However, there is yet no evidence for any phenotypic relevance of copy number variation in these genes (see also above). Thus, we conclude that biosynthesis of the two major cannabinoids THCA and CBDA are the result of presence/absence, sequence variation, and expression of single copy genes (McKernan et al. 2020).

### Gene sequence variation and potential geographic origins

Besides the divergence at the genomic level mentioned above, sequence variation within cannabinoid oxidocyclase gene sequences may help shed light on their evolutionary history. For example, subclade A4 pseudogenes are generally single copy but can be divided into two divergent types (Table 1; Figs 3, S1, S2). Type 1 can be found in CBDA-dominant cultivars ‘Finola’, ‘CBDRx’, and on the *CBDAS*-containing haplotype of cultivar ‘Jamaican Lion’. Type 2 on the other hand can be found in drug-type cultivars ‘Purple Kush’, ‘LA confidential’, and on the *THCAS*-containing haplotype of cultivar ‘Jamaican Lion’. Similarly, the full-length subclade A3 gene that is closely linked to *CBDAS* in the genome of cultivar ‘Finola’ is sister to the subclade A3 pseudogenes closely linked to *THCAS* in cultivars ‘Purple Kush’, ‘Jamaican Lion’, and the plant from Jilong (Figure 3). These findings further corroborate our evolutionary interpretation of cannabinoid oxidocyclase gene duplication and divergence shown in Figure 5 and suggest significant and consistent divergence between haplotypes containing *CBDAS* and *THCAS*. Thus, genomic divergence described above correlates with prevalence of THCA and CBDA and, hence, perhaps with genetic origins of drug-versus fiber type cultivars. Drug-type cultivars are considered to have originated in two different regions of the Himalayan foothills, while fiber-type cultivars are considered to have been developed independently in Europe and in East Asia (Clarke and Merlin 2013; Clarke and Merlin 2016). Thus the observed genetic variation may be a consequence of divergence between these different geographic regions. Similarly, variation within *THCAS* sequences may also reflect geographic origin. Level of sequence divergence between *THCAS* from different geographic areas in South Korea is relatively high (Table S3, Figure S1). Accessions from Boseung province share three unique aa substitutions, the accession from Jecheon province in Korea that has five unique aa substitutions, and sequences from Cheungsam share ten unique aa substitutions (Doh et al. 2019). Moreover, the *THCAS* sequence from the putatively wild accession from Jilong, Tibet is also relatively different from canonical THCAS and placed as sister to all other clade A1 sequences (S. Gao et al. 2020). This suggests that additional sampling throughout the native range of *Cannabis* is likely to reveal new genetic variation. However, germplasm from regions of origin is scarce, especially when restricting samples to those that are compliant with international regulations such as the Nagoya-protocol. We therefore strongly support earlier calls for increased efforts to develop well-curated public germplasm banks covering *Cannabis’* entire natural variation (Welling et al. 2016; Small 2018; Kovalchuk et al. 2020; McPartland and Small 2020).

**Table 1.** Number of full-length coding and nonfunctional cannabinoid oxidocyclase sequences in high-quality genome assemblies.

|  | Finola |  | Purple Kush |  | CBDRx |  | Jamaican Lion |  | Jilong |  |
| --- | --- | --- | --- | --- | --- | --- | --- | --- | --- | --- |
| Clade | coding | nonfunctional | coding | nonfunctional | coding | nonfunctional | coding | nonfunctional | coding | nonfunctional |
| A1_THCAS | 0 | 0 | 1 | 0 | 0 | 0 | 1 | 0 | 1 | 0 |
| A2_CBCAS | 2 | 4 | 2 | 3 | 1 | 5 | 6 | 2 | 0 | 0 |
| A3 | 1 | 0 | 0 | 1 | 0 | 0 | 0 | 1 | 0 | 2 |
| A4 | 0 | 1 | 0 | 1 | 0 | 1 | 0 | 2 | 0 | 0 |
| B1_CBDAS | 1 | 0 | 0 | 0 | 1 | 0 | 1 | 0 | 0 | 0 |
| B2 | 0 | 0 | 0 | 3 | 0 | 0 | 0 | 3 | 0 | 5 |
| C | 0 | 5 | 0 | 4 | 1 | 4 | 7 | 3 | 0 | 0 |
| sums | 4 | 10 | 3 | 12 | 3 | 10 | 15 | 11 | 1 | 7 |
| total | 14 |  | 15 |  | 13 |  | 26 |  | 8 |  |

## Materials & Methods

### Sequence sampling

The sequence data underlying this article are available in the GenBank Nucleotide Database at www.ncbi.nlm.nih.gov, and can be accessed with the accession codes listed in Tables S2-S4.

#### Sampling of berberine bridge protein sequences

We sampled full-length predicted berberine bridge protein sequences from the Eurosid clade based on whole-genome assemblies of *Cannabis sativa* cultivar ‘CBDRx’ (GCF_900626175.1), *Humulus japonicus* cultivar ‘Cascade’ (from http://hopbase.cgrb.oregonstate.edu), *Parasponia andersonii* (GCA_002914805.1), *Trema orientale* (GCA_002914845.1), *Morus notabilis* (GCF_000414095.1), *Medicago truncatula* (JCVI MedtrA17_4.0), and *Arabidopsis thaliana* (TAIR10) (Tang et al. 2014; Berardini et al. 2015; Grassa et al. 2018; van Velzen et al. 2018). Some *Cannabis* and *Humulus* genes were found to be misannotated or lacking an annotation. These were manually corrected based on alignment with a closely related and correctly annotated genome sequence. Because the CBDRx genome does not include *THCAS*, we included accession Q8GTB6.1 (Sirikantaramas et al. 2004). We also indicated the putative orthologs of *Morus alba* Diels–Alderase (MaDA) and moracin C oxidase (MaMO) in *Morus notabilis* and included daurichromenic acid synthase from *Rhododendron dauricum* (accession BAZ95780.1) and berberine bridge enzyme from *Eschscholzia californica* (accession AAC39358.1) (Hauschild et al. 1998; Iijima et al. 2017; L. Gao et al. 2020). Sequences of *Arabidopsis thaliana* CYTOKININ OXIDASE 1 and 2 were used as outgroups.

#### Sampling of cannabinoid oxidocyclase nucleotide sequences

whole-genome assemblies of cultivars We mined available near chromosome-level genome assemblies of *Cannabis sativa* cultivars ‘CBDRx’ (GCF_900626175.1), ‘Jamaican Lion’ (mother: GCA_012923435.1) (McKernan et al. 2018; McKernan et al. 2020), ‘Finola’ (GCA_003417725.2), ‘Purple Kush’ (GCA_000230575.4) (van Bakel et al. 2011; Laverty et al. 2019), and a putatively wild plant from from Jilong, Tibet (GCA_013030365.1) (S. Gao et al. 2020) for homologs of characterized cannabinoid oxidocyclase sequences (i.e. *THCAS, CBDAS, CBCAS*) using BLASTP and BLASTN implemented in Geneious Prime 2019 (https://www.geneious.com). We similarly mined sequences from additional genome assemblies of cultivars ‘Cannatonic’ (GCA_001865755.1), ‘Chemdog91’ (GCA_001509995.1), ‘Jamaican Lion’ (father) (GCA_013030025.1), ‘LA confidential’ (GCA_001510005.1), and ‘Pineapple Banana Bubble Kush’ (GCA_002090435.1) (McKernan et al. 2018; Vergara et al. 2019; McKernan et al. 2020). When necessary, structural annotations were manually modified based on nucleotide alignments with annotated genes with highest identity. When genes comprised putative pseudogenes (i.e. coding sequence was fragmented due to premature stop codons and/or frame-shifts) they were annotated manually such that CDS remained homologous and all nonsense and frameshift mutations were indicated.

In addition, we compiled homologous nucleotide sequences available from ncbi databases the majority of which came from published studies (Sirikantaramas et al. 2004; Kojoma et al. 2006; Taura, Sirikantaramas, Shoyama, Yoshikai, et al. 2007; El Alaoui et al. 2013; McKernan et al. 2015; Onofri et al. 2015; Weiblen et al. 2015; Cascini et al. 2019; Doh et al. 2019). Based on preliminary analyses, some sequences described by (Cascini et al. 2019) were found to have relatively high levels of ambiguous nucleotides, probably due to unspecific amplification of multiple genes or gene variants. Some sequences amplified from Moroccan hashish samples described by (El Alaoui et al. 2013) were suspected to be chimeric (probably due to differential specificity between forward and reverse sequencing primers). These ambiguous and suspected chimeric sequences were excluded from final analyses. The most closely related BBE-like sequences from *Cannabis* and *Humulus* were used as outgroups.

### Phylogenetic analyses

Multiple sequence alignment was performed with MAFFT v7.450 with automatic selection of appropriate algorithm, a gap open penalty of 1.26 and an offset value 0.123. For protein and nucleotide sequence datasets we used the BLOSUM62 and 100PAM scoring matrix, respectively. Optimal models of sequence evolution as determined using Modeltest-NG v.0.1.5 on XSEDE via the CIPRES gateway (Miller et al. 2010; Darriba et al. 2020) were WAG+I+G4 for the protein data set and GTR+I+G4 for all nucleotide data sets. Gene trees were reconstructed in a Bayesian framework using MrBayes v 3.2.6 (Ronquist and Huelsenbeck 2003) implemented in Geneious Prime with a chain length of 2.2 million generations; sampling every 1000th generation; 4 heated chains with a temperature of 0.2. The first 200,000 generations were discarded as burnin.

Within the berberine bridge gene family tree, clades were numbered in accordance with earlier classification (Daniel et al. 2016). Within the cannabinoid oxidocyclase gene tree, clades and types were characterized based on unique nonsynonymous substitutions (i.e. substitutions resulting in a change to a specific amino acid that, based on current sampling, were unique for as well as constant within that clade) where possible. All site numbers were projected to the THCAS reference sequence described by (Sirikantaramas et al. 2004) (accession AB057805.1) and, within the THCAS clade, type names were kept in accordance with those from Onofri et al. (Onofri et al. 2015).

The multiple-sequence alignments and associated gene trees are available in the 4TU.ResearchData repository at https://doi.org/10.4121/13414694.

### Microsynteny assessment

Nucleotide microsynteny was assessed for the cultivars ‘Jamaican Lion’ (mother), ‘CBDRx’, ‘Finola’, ‘Purple Kush’, and a putatively wild specimen from Jilong, Tibet. Because we found inconsistencies between the different genome assemblies in the ordering and orientation of sequences into scaffolds we considered genomic contigs only. Nucleotide-level alignments were generated by performing gapped extensions of high-scoring segment pairs using Lastz version 1.04.03. To avoid seeding in repetitive sequence we indexed unique words with single alignments only (--maxwordcount=1; --masking=1). To reduce runtime we set --notransition and --step=20. To keep tandem repeats we set --nochain. To filter short and/or dissimilar alignments we set --hspthresh=100000., --filter=identity:95, and --filter=nmatch:2000. The --rdotplot output was used to generate alignment dotplots in R. For nucleotide level microsyntenic blocks of interest we further assessed microsynteny with genomic sequences from *H. lupulus* cultivar ‘Cascade’, *P. andersonii*, and *T. orientale* based on predicted protein sequences (van Velzen et al. 2018; Padgitt-Cobb et al. 2019).

## Acknowledgements

We thank Bob Harris (Pennsylvania State University, U.S.A.) for advice on generating dotplots based on Lastz output. Marleen Botermans (NVWA, The Netherlands) provided valuable comments on an early draft of the manuscript. Bastian Daniel (Graz University, Austria) kindly helped with BBE-like gene family classification.

## Supplementary tables

Table S1. Clade, subclade, and type-specific unique nonsynonymous substitutions and nonsense mutations. Column “Onofri type” lists type numbers as defined by Onofri et al. (2015); n.d.: no difference.

Table S2. Sampling of berberine bridge enzyme protein sequences.

Table S3. Sampling of cannabinoid oxidocyclase nucleotide amplicon sequences

Table S4. Sampling of cannabinoid oxidocyclase nucleotide sequences based on genomic contigs. Asterisks indicate manually corrected gene models.

## Supplementary figure legends

Fig. S1. Cannabinoid oxidocyclase gene tree based on nucleotide sequences from genbank accessions. Clade B was used as an outgroup. Labels indicate genbank accession; putative nonfunctional (pseudo)genes are in grey; functionally characterized THCAS and CBDAS (Sirikantaramas et al. 2004; Taura, Sirikantaramas, Shoyama, Yoshikai, et al. 2007) are indicated in dark red. Coloured blocks indicate the identified clades; white blocks indicate sequence types. Node labels indicate posterior probabilities below 1.0.

Fig. S2. Cannabinoid oxidocyclase gene tree based on nucleotide sequences from whole-genome assemblies of cultivars ‘Cannatonic’, ‘Chemdog91’, ‘Jamaican Lion’ (father), ‘LA confidential’, and ‘Pineapple Banana Bubble Kush’ (PBBK). Clade B was used as outgroup. Labels indicate genbank accession of genomic contig and locus tag (when available) or start position. Putative nonfunctional (pseudo)genes are in grey; functionally characterized THCAS, CBDAS, and CBCAS (Sirikantaramas et al. 2004; Taura, Sirikantaramas, Shoyama, Yoshikai, et al. 2007; Laverty et al. 2019) are indicated in dark red. Coloured blocks indicate the identified clades; white blocks indicate sequence types. Node labels indicate posterior probabilities below 1.0.

Fig. S3. LASTZ nucleotide alignment dotplots of microsyntenic cluster 1 showing clade C tandemly repeated array and (A) CBDAS, or (B) THCAS variants. Sense alignments are in black; antisense alignments are in blue. Triangles indicate start positions of genes. For gene color codes see Fig. 4A.

Fig. S4. LASTZ nucleotide alignment dotplots of microsyntenic cluster 2 showing CBCAS tandemly repeated array. Sense alignments are in black; antisense alignments are in blue. Triangles indicate start positions of genes. For gene color codes see Fig. 4B.

