## Supplementary figures and images for "Origin and evolution of the cannabinoid oxidocyclase gene family"

### Supplemental Figure S1

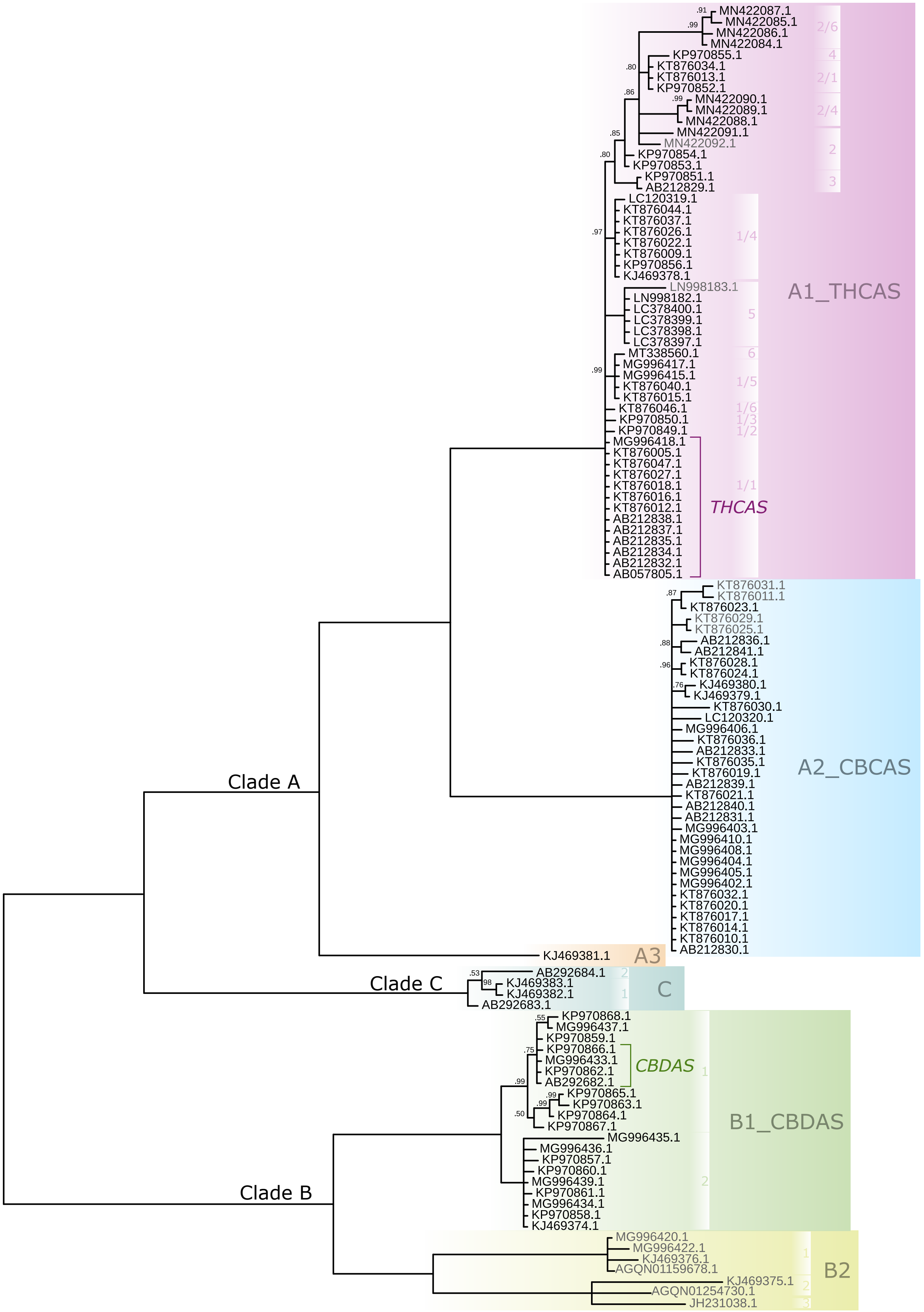

### Supplemental Figure S2

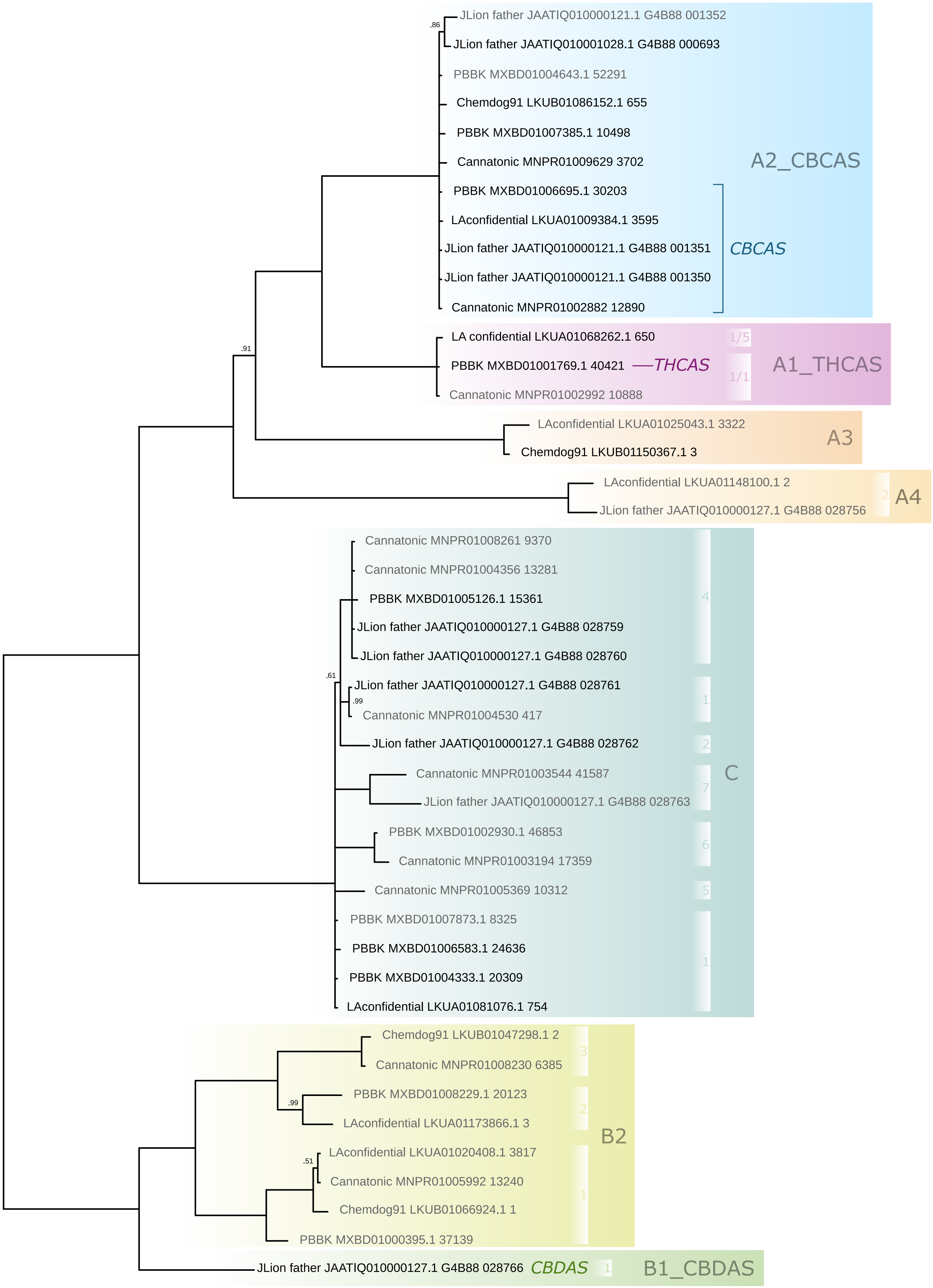

### Supplemental Figure S3

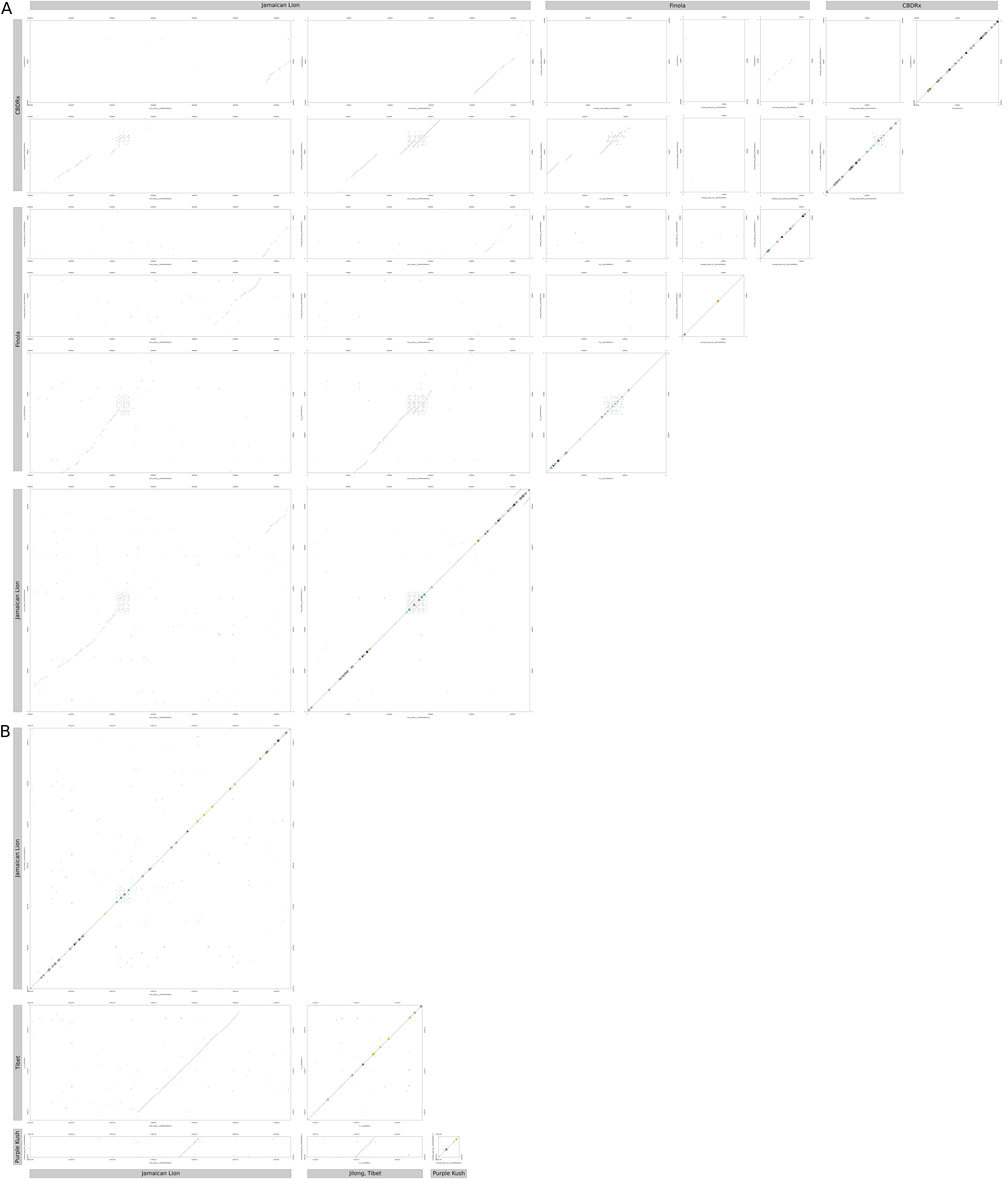

### Supplemental Figure S4

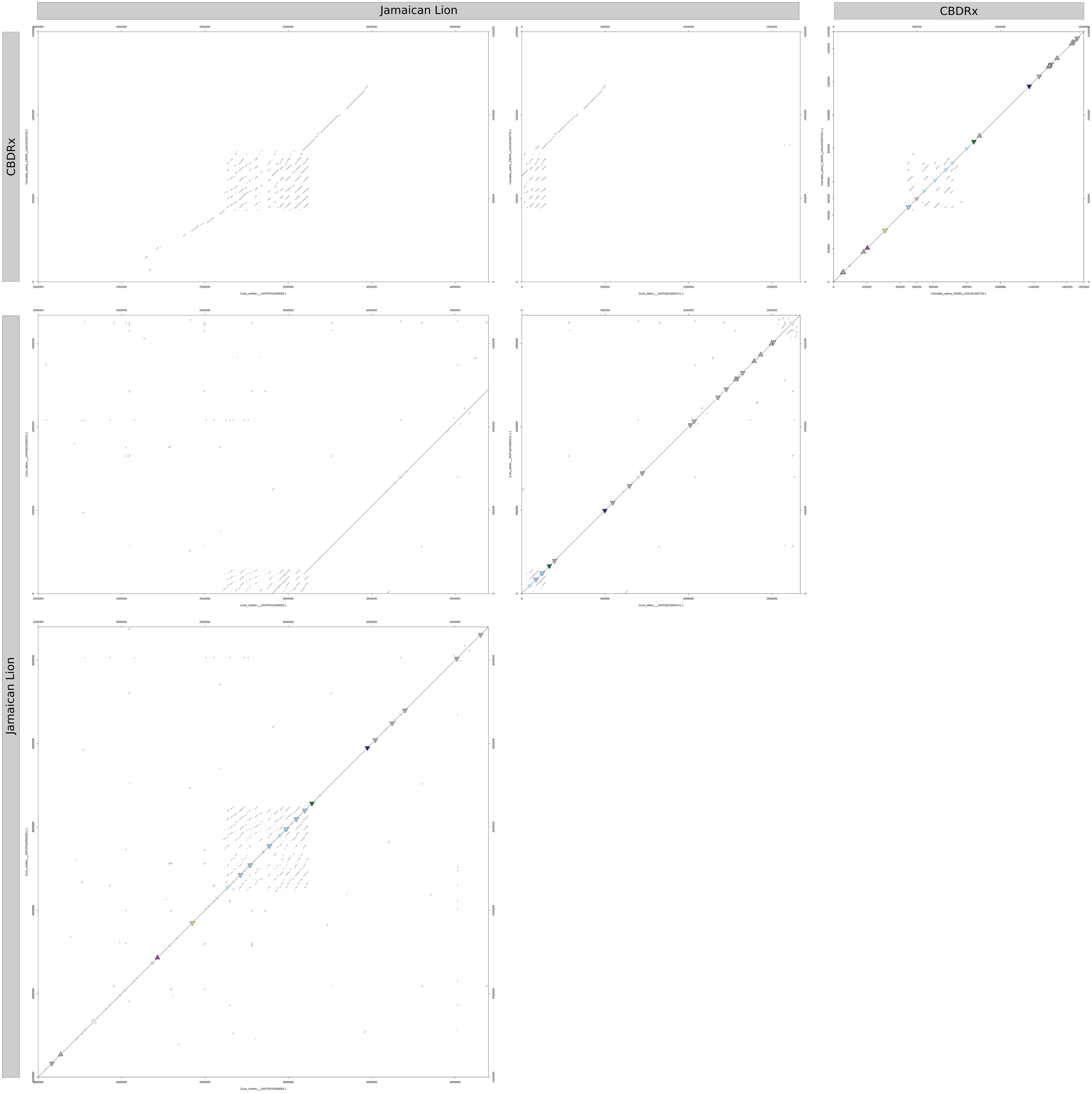
